# *Pseudomonas aeruginosa* Intra-Population Diversity Shapes Host Airway Epithelial Responses

**DOI:** 10.64898/2026.08.11.744332

**Authors:** Stephen P. Diggle, Sheyda Azimi

**Affiliations:** School of Biological Sciences, Georgia Institute of Technology, Atlanta, GA, USA; Department of Biology, College of Arts and Sciences, Georgia State University, Atlanta, GA, USA

**Keywords:** *Pseudomonas aeruginosa*, cystic fibrosis, within-host diversity, intra-population interactions, inflammation, transcriptomics, chronic infection

## Abstract

In chronic pulmonary infection airways are colonized by genetically and phenotypically diverse *Pseudomonas aeruginosa* populations, yet almost everything known about *P. aeruginosa* pathogenesis has been learned from single clinical isolates or laboratory reference strains studied in isolation. Whether a diverse population behaves as the sum of its members remains unclear. We infected differentiated primary cystic fibrosis (CF) airway epithelial cells (CF-pAECs) at air-liquid interface with whole *P. aeruginosa* populations collected from the sputum of three adults with CF, and, in parallel, with genetic variants retrieved from the same populations. We found that host responses to whole populations were not predicted by responses to their derivative variants, and mixed populations did not exhibit the average expected response of the members. Variation in host response was dominated by the tissue-remodeling mediators vascular endothelial growth factor (VEGF) and matrix metalloproteinase-9 (MMP-9), which differed distinctly between individual variants, whereas core pro-inflammatory cytokines (IL-6, IL-8, TNF-α, IFN-γ) varied comparatively little. Bacterial transcriptomes recorded during infection showed the same asymmetry. Clinical populations shared a transcriptional state distinct from PAO1, and mixed populations maintained stable expression of core regulatory, secretion, and DNA-repair loci (including *hfq*, *xcpT*, and *ssb*), while the derivative variants grown alone exhibited differential transcriptional profiles. Interactions among co-existing lineages therefore shape both bacterial physiology and host response, suggesting that the diverse population, not the single clone, is the appropriate unit of study in chronic infection.

**Importance:** In chronic respiratory infections, lungs are not colonized by a clonal population but by diverse co-evolving *Pseudomonas aeruginosa* lineages that have often evolved together for many years. By comparing primary human cystic fibrosis (CF) airway epithelial cells responses to whole *P. aeruginosa* populations taken from sputum against their isolated sub-lineages, we demonstrate that host inflammatory responses to the population could not be predicted from the responses to their derivative individual variants, and the bacteria themselves behaved differently in a population than they did alone. These findings highlight that working on single isolates risk misrepresenting chronic infection biology and emphasize that effective therapeutic strategies must target population-level dynamics rather than single isolated strains.

## Introduction

Bacteria associated with humans live in diverse microbial communities, where their physiology is shaped both by host responses and by neighboring bacterial cells (1). In chronic infection, within-host evolution generates co-existing lineages of the same species (and often strain) that differ in antimicrobial resistance and virulence-associated traits (2–4). How this intra-population structure alters the course of infection is a central open question in understanding the mechanisms of pathogenesis of chronic infections.

Chronic respiratory infection with *Pseudomonas aeruginosa* remains a major cause of declining lung function and bronchiectasis in people with cystic fibrosis (CF) and chronic obstructive pulmonary disease (COPD) (5–7). In CF, dysfunction of the CF transmembrane conductance regulator (CFTR) dehydrates the airway surface liquid and impairs mucociliary clearance, producing hypoxia, epithelial damage, and a chronic inflammatory state, driving the airway injury, thus epithelium responses determine the damage attributed to chronic infection (8–12). A hallmark of chronic respiratory infections with *P. aeruginosa* is intra-population diversity, where pathoadaptive mutations accumulate, and multiple genetic variants co-occur within a single sputum sample at any one time (13, 14). Mutations in genes encoding quorum sensing (QS) systems, motility, antimicrobial resistance (AMR), lipopolysaccharide (LPS), and O-antigen are common (14–19). Intra-population diversity itself has been associated with reduced lung function (20, 21), but the mechanisms linking *P. aeruginosa* population diversity to airway damage remain unresolved.

We have previously shown that allelic polymorphism and changes in population structure modulate collective functional phenotypes such as QS, protease production and AMR in evolving populations (22), that population-level genetic diversity does not predict the intrastrain AMR heterogeneity contained within a population (23), and that within-population polymorphism shapes airway microbiogeography at the micron scale (24, 25). What has not been tested is whether the host response to a diverse *P. aeruginosa* population can be predicted from the immune responses to the genetic variants that compose it. This question becomes important as *P. aeruginosa* transcriptional signatures recovered from expectorated sputum samples or explanted CF lungs, display similar transcriptomic profiles which are broadly conserved across patients (26–28), suggesting that population structure is transcriptionally inconsequential, and that a single isolate can be an adequate proxy for the population.

Here we tested whether intra-population diversity changes bacterial physiology and host responses during infection of human airway epithelium. We infected differentiated primary CF airway epithelial cells (CF-pAECs) at air–liquid interface (ALI) with whole *P. aeruginosa* populations from three adults with CF with chronic infection but differing lung function, and, in parallel, with genetic variants from the same populations, and we measured host immune responses and *P. aeruginosa* transcriptional profiles. We show that (i) the epithelial response to mixed *P. aeruginosa* populations is not predicted by the responses to single variants; (ii) difference in epithelial response is dominated by tissue-remodeling mediators rather than by core pro-inflammatory cytokines; and (iii) mixed populations maintained a stable expression of core regulatory, secretion, and DNA-repair genes that diverges in the single variants grown alone. Together, our data demonstrate that the population, and not single isolates, serves as the unique functional unit in chronic *P. aeruginosa* infection depending on spatial segregation across localized airway niches. Considering the focal injury patterns typical of bronchiectasis, determining how micron-scale spatial partitioning and population structure interact locally remains vital to understanding host-pathogen dynamics in chronic respiratory infections.

## Results

### Collection of whole *P. aeruginosa* populations and their constituent variants from adults with CF

We collected expectorated sputum from three adults with CF with chronic *P. aeruginosa* infection and differing lung function (Fig. 1a and b). Diluted sputum was plated on *Pseudomonas* isolation agar (PIA), and all grown colonies were collected and stored together as whole mixed populations (P1, P2, P3). To capture intra-population phenotypic diversity, we plated subcultures on Congo red agar and archived a single representative colony of each distinct morphotype as a clonal variant. We archived four variants from P1 (P1V1–P1V4), three from P2 (P2V1–P2V3), and one from P3 (P3V1) (Fig. 1c).

**Fig. 1.**
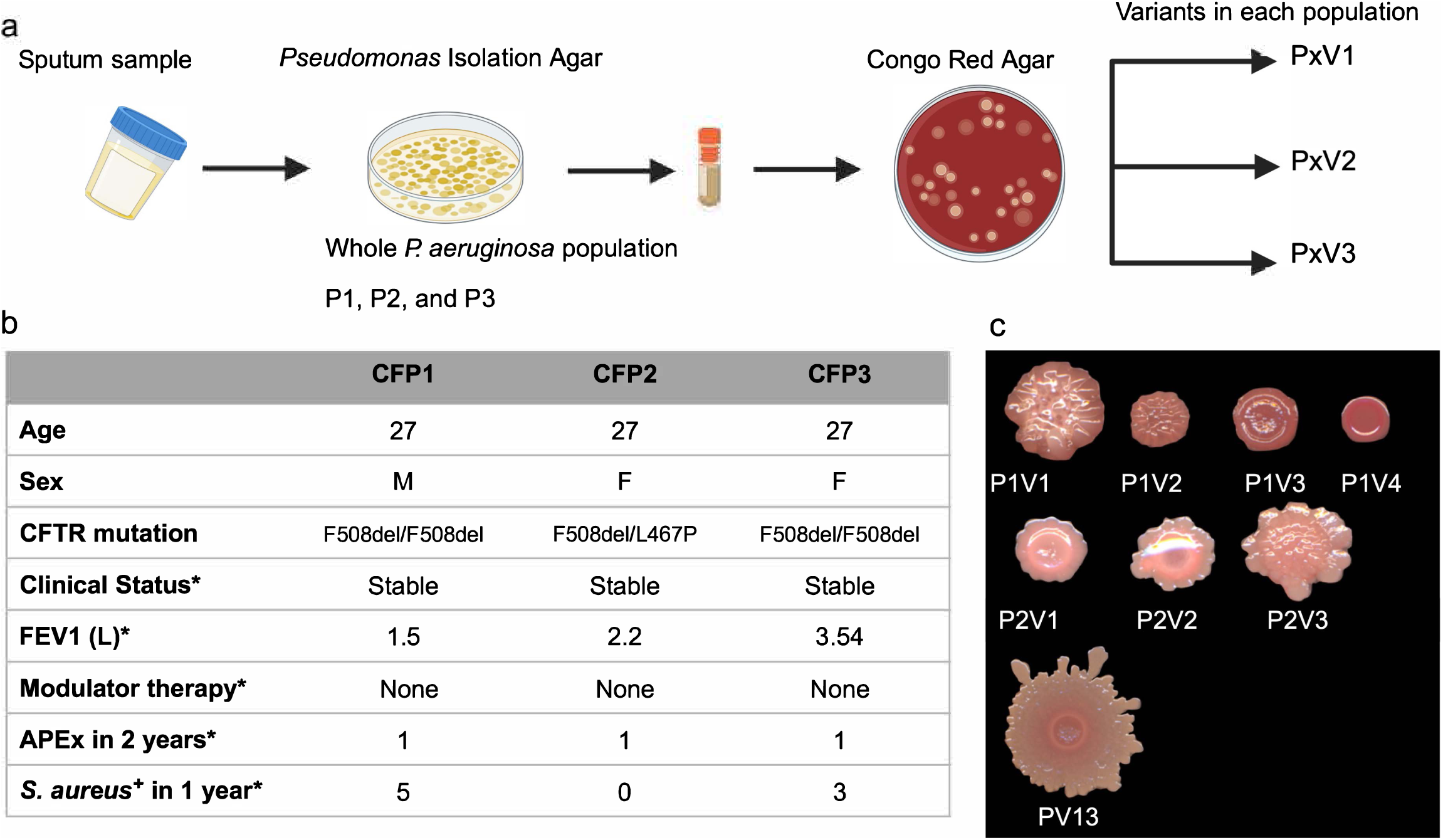
Participant characteristics and colony morphology of clinical *P. aeruginosa* populations and their clonal variants. (a) Schematic of the collection of whole mixed *P. aeruginosa* populations and clonal variants. Sputum was plated on *Pseudomonas* isolation agar to collect whole populations (P1, P2, P3), and subcultures were plated on Congo red agar to identify and isolate phenotypically distinct variants from each population. (b) Characteristics of participants (CFP1, CFP2, CFP3), including CFTR modulator therapy at the time of sputum collection, frequency of acute pulmonary exacerbations in the preceding 2 years (APEx), and positive *Staphylococcus aureus* culture within the year before sputum collection. (c) Colony morphology of individual variants isolated from P1 (P1V1–P1V4), P2 (P2V1–P2V3), and P3 (P3V1).

### The airway epithelial response to a whole population is not predicted by its constituent variants

Bacterial pathogenesis and host immune activation are conventionally assessed using isogenic mutants or single clinical isolates, hence the influence of intra-population diversity on host responses is largely untested. To address this, we infected differentiated CF-pAECs at ALI with either whole *P. aeruginosa* populations or their individual variants (≈10^3^ CFU in 100 µL SCFM2 per transwell; MOI ≈ 0.01) for 12 h, and quantified secreted cytokines and chemokines in the basolateral compartment. Populations from each CF participant induced distinct inflammatory profiles (Fig. S1a–c). Critically, in principal component analysis (PCA) of the full cytokine panel, whole populations did not display the average predicted value of the inflammatory responses of their own constituent variants (Fig. 2a; PC1 43.12%, PC2 39.08%, 82.20% of total variance). Variants from P2 and P3 clustered tightly, whereas variants from P1 separated into distinct regions, driven principally by the outlying variant P1V1 along PC1. The immune response to each mixed population was therefore not an average of the inflammatory responses of its members, indicating that interactions among co-existing variants generate host signaling outcomes that are not predictable from the single variants.

**Fig. 2.**
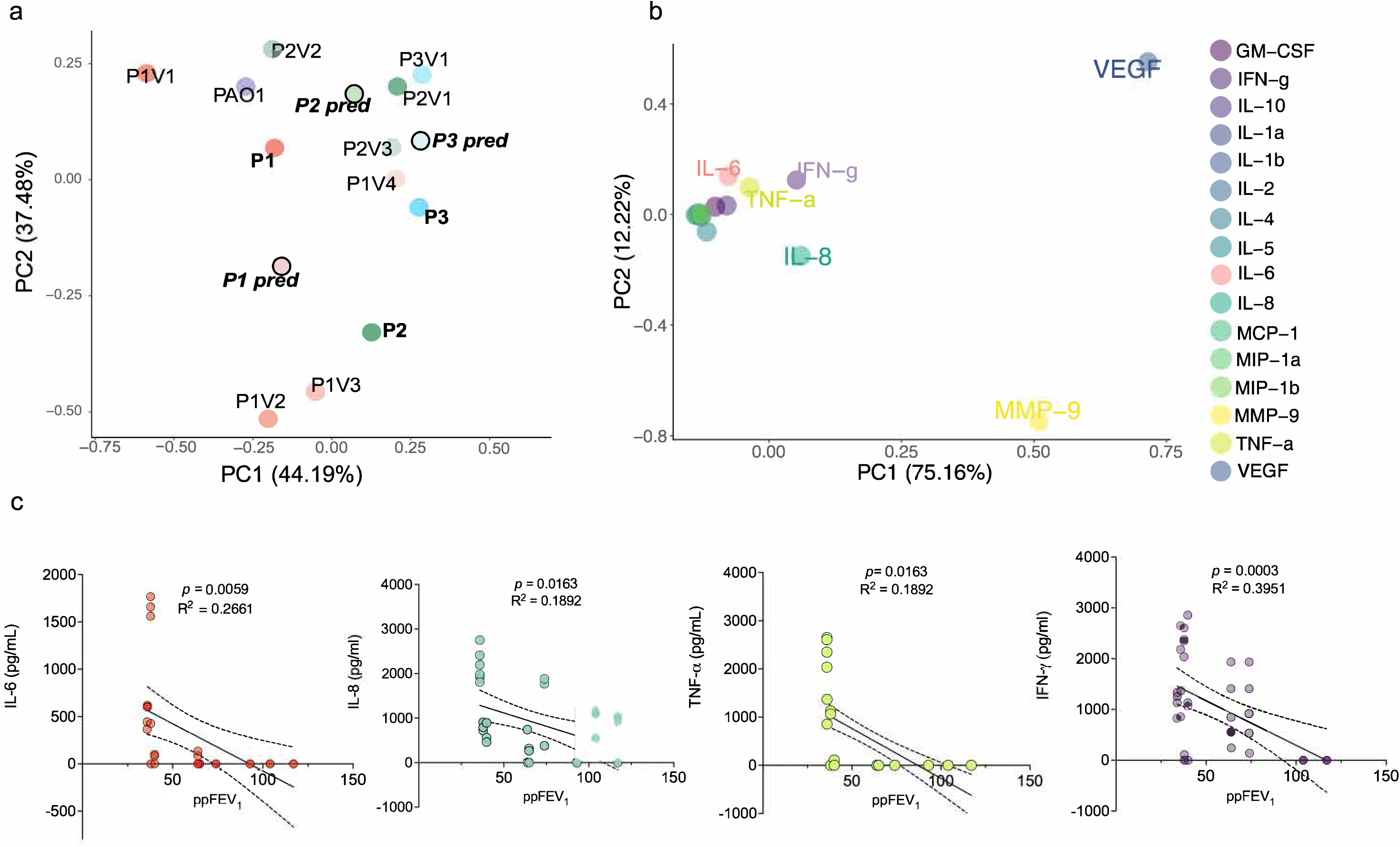
The epithelial response to a whole *P. aeruginosa* population is not predicted by its constituent variants. (a) PCA of cytokine levels produced by CF-pAECs infected with clinical variants, whole mixed populations, and PAO1. PC1 (44.19%) and PC2 (37.48%) explain 81.67% of the total variance in host response. Symbols are colored by participant of origin (P1, red; P2, green; P3, blue; PAO1, purple). Open circles with bold black borders represent the predicted additive population centroids (P1 pred, P2 pred, P3 pred) calculated by averaging the constituent sub-clonal variant responses for each patient population with distinct separation between observed populations and their corresponding predicted additive centroids. Variants from P2 and P3 cluster tightly, whereas variants from P1 separate into distinct regions, driven by the outlying variant P1V1 along PC1. (b) PC1 (75.16%) and PC2 (12.22%) explain 87.38% of the variance in cytokine production. Core pro-inflammatory cytokines (IL-6, IL-8, TNF-α, IFN-γ) cluster together, while the tissue-remodeling mediators VEGF and MMP-9 separate along PC1. (c) Linear regression analysis comparing the participant ppFEV_1_ against core pro-inflammatory cytokines display a significant negative correlation with clinical lung function. (IL-6 (*R*^2^ = 0.2661, *P* = 0.0059), IL-8 (*R*^2^ = 0.1892, *P* = 0.0163), TNF-α (*R*^2^ = 0.1892, *P* = 0.0163), and IFN-γ (*R*^2^ = 0.3951, *P* = 0.0003) Solid lines show best-fit linear regressions; dashed curves show 95% confidence intervals).

### Variation in host output is dominated by tissue-remodeling signals rather than core cytokines

To identify the cytokines driving the difference in inflammatory between infection conditions, we performed PCA on the measured cytokines themselves. Variation was dominated by the tissue-remodeling signals, vascular endothelial growth factor (VEGF) and matrix metalloproteinase-9 (MMP-9) (Fig. 2b; PC1 75.16%), while the core pro-inflammatory cytokines IL-6, IL-8, TNF-α, and IFN-γ, which clustered together, accounted for far less of the variance (PC2 12.22%) in epithelial responses to *P. aeruginosa* infection. Individual variants differed markedly in the levels of VEGF and MMP-9 they elicited (Fig. 2b, Fig. S1), whereas the core cytokine response was comparatively uniform across *P. aeruginosa* populations or variants. Different variants from the same sputum sample/airway therefore do not simply elicit different inflammation levels; they shift the epithelium between qualitatively different pathways, with some driving a remodeling response rather than a conventional acute antibacterial one.

Consistent with the reported association between within-host *P. aeruginosa* diversity and disease severity, IL-6, IL-8, TNF-α, and IFN-γ levels negatively associated with each participant baseline lung function (ppFEV_1_) (Fig. 2c) (20, 21), while there was no such association for VEGF and MMP-9 levels (Fig. S1d and e), reinforcing that the remodeling axis and the acute cytokine axis are independently regulated by bacterial input. while our analysis derived from only three participants, it shows a link between differential immune responses by airway epithelial cells and loss of lung function.

### Clinical populations exhibit distinct transcriptional profiles during epithelial infection

To ask whether the divergent host responses were accompanied by divergent bacterial physiology, we performed bulk RNA sequencing on *P. aeruginosa* recovered from the same infections. Global transcriptional profiles separated by participant of origin, with PC1 and PC2 accounting for 58% of expression variance (40% and 18%, respectively) (Fig. 3a). Variants from P2 are transcriptionally distinct from the other clinical samples and from PAO1; variants derived from P1 show distinct transcriptomes from the mixed P1 parental population along PC2; while P1, P3, P3V1, and PAO1 display similar transcriptional profiles. Using variance partitioning analysis, we identified 47 genes whose expression separated clinical populations from PAO1 as two opposing blocks (Fig. 3b and c, Table S1). In PAO1 *PA4686*, *PA2208*, *moaB2*, and *PA2453* were highly expressed and downregulated across all clinical populations, whereas *PA4089*, *pauA6*, *PA2707*, and the SOS-induced cell division inhibitor *sulA* were consistently upregulated in the clinical populations.

**Fig. 3.**
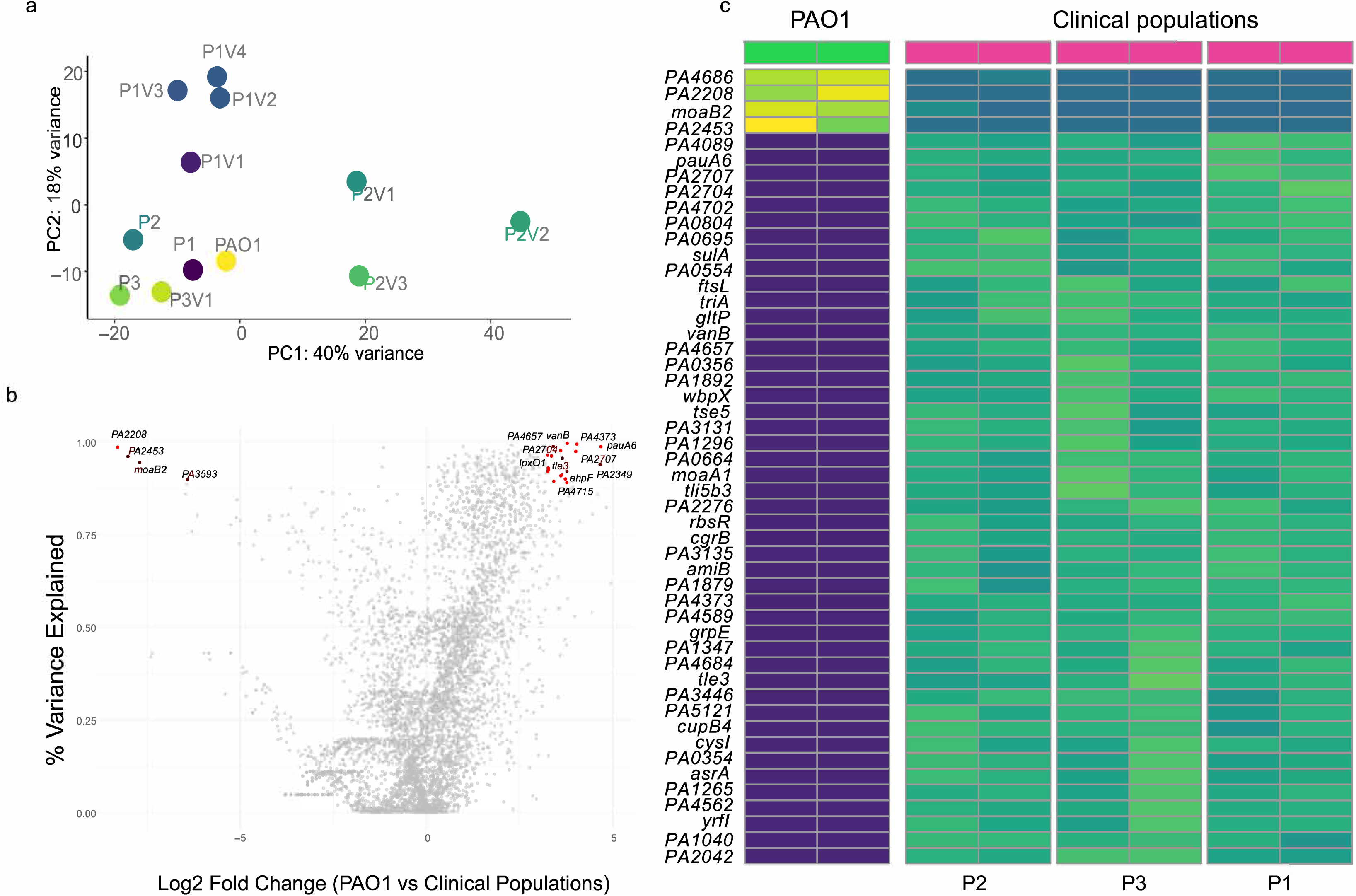
Clinical *P. aeruginosa* populations occupy a transcriptional state distinct from PAO1 during epithelial infection. (a) Global transcriptional profiles of mixed populations (P1, P2, P3), their derivative variants, and PAO1. PC1 and PC2 capture 58% of total transcriptional variance, with mixed populations and variants derived from P2 (P2V1–P2V3) separating along PC1 and variants derived from P1 clustering along PC2. (b) Genes identified by variance partitioning as consistently differentially expressed between clinical populations and PAO1. Highlighted red loci represent top driver transcripts variance is only present in clinical populations (>90%), separating PAO1-enriched baseline transcripts (e.g., *PA2208*, *PA2453*, *moaB2*) from upregulated clinical markers (e.g., *pauA6*, *vanB*, *PA2707*, *PA4657*). (c) Expression of the 47 genes identified by variance partitioning, driving the opposite expression levels of *PA4686*, *PA2208*, *moaB2*, and *PA2453* by PAO1 and across clinical populations, while *PA4089*, *pauA6*, *PA2707*, *sulA*, *PA1040*, and *PA2042* are stably upregulated across all clinical populations.

### Mixed populations stabilize expression of core regulatory genes but diverge in individual variants

Additional PCA analysis showed that three mixed populations were transcriptionally distinct (Fig. 4a; PC1 49%, PC2 30%). We found that expression of genes in oxidative phosphorylation and nitrogen metabolism pathways by P1 and P2 physiology, and differential expression of genes involved in biofilm formation by P3 are key distinguishing features between these populations (Fig. 4c, Fig. S2, Table S2).

**Fig. 4.**
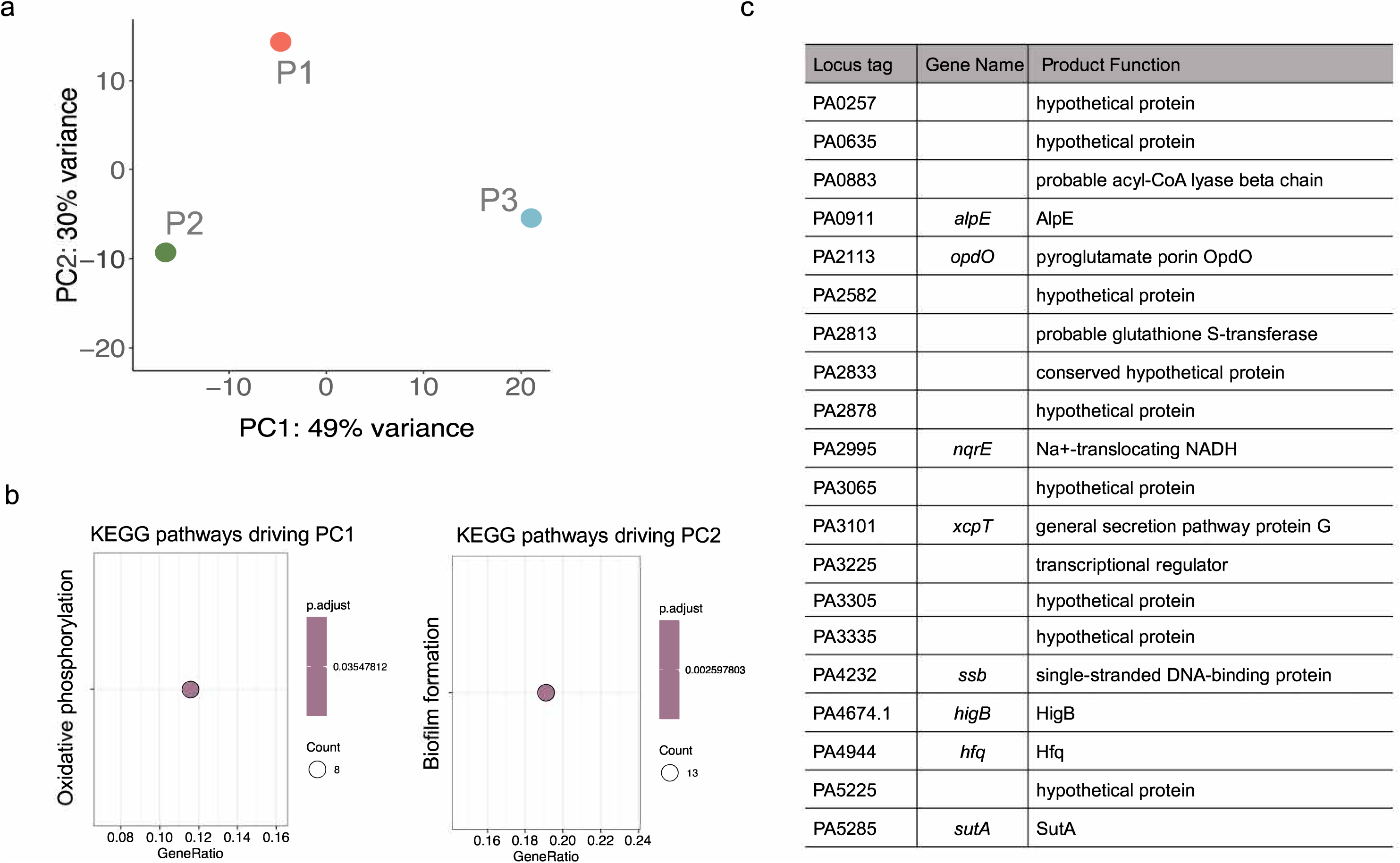
Mixed populations keep a stable gene expression for core regulatory, secretion, and DNA-repair system. (a) Global transcriptomic separation among mixed populations (P1, P2, P3). PC1 (49%) and PC2 (30%) capture 79% of total transcriptional variance, with P3 separating along PC1 and P1 along PC2 relative to P2. (b) Driver loci identified by variance partitioning that separate population-level transcriptomes from those of individual variants, including *hfq* (PA4944), *xcpT* (PA3101), and *ssb* (PA4232). (c) KEGG/GO enrichment of genes driving variation between clinical populations, showing enrichment of oxidative phosphorylation and nitrogen metabolism along PC1 and biofilm formation along PC2.

To identify the transcriptional signatures that distinguish a population from its own members, we analyzed the top transcripts from variance partitioning model. This revealed a set of genes with variable expression in the mixed-population group, with negligible intergroup variance (<0.01%). These loci include the post-transcriptional regulator *hfq* (PA4944), the type II secretion system component *xcpT* (PA3101), and the single-stranded DNA-binding protein *ssb* (PA4232) (Fig. 4b). Direct comparison of each variant with its parental population (Fig. S3–S5, Table S2) showed the complementary pattern, where in individual variants genes involved in adhesion, phenazine biosynthesis, and secondary metabolism including the *cupE* fimbrial cluster (*cupE5*, *cupE6*) and *phzE1* were downregulated relative to their parental populations. while mixed parental populations maintained a stable expression across the core regulatory, secretion, and DNA-repair functions.

## Discussion

Understanding how intra-host bacterial diversity shapes infection dynamics remains a challenge in chronic infection. *P. aeruginosa* pathogenesis is almost always studied by using single clinical isolates or laboratory reference strains. However, chronic pulmonary infections such as CF and Non-CF bronchiectasis are dominated by complex, heterogeneous bacterial populations rather than clonal populations. In this study, we demonstrate that inflammatory responses of CF-pAECs to diverse *P. aeruginosa* population could not be predicted from the responses to their constituent sub-lineages. Additionally, bacteria exhibited different physiological states as mixed diverse populations when compared with individual isolates sampled from those populations. Therefore, the behavior of both host and pathogen are altered due to *P. aeruginosa* intra-population heterogeneity.

Because strain background, growth media, and infection conditions were strictly controlled, the observed non-additivity of the host epithelial response represents an intrinsic property of community behavior. The most compelling explanations for this uncoupling are functional interactions that emerge exclusively within heterogeneous populations, such as metabolic cross-feeding, the cooperative exchange or exploitation of secreted public goods, and physical inter-lineage aggregation that alters the structural interface presented to host epithelial cells (19, 29). While delineating the precise molecular drivers of this non-additivity was beyond the scope of the present study, systematically decoupling these inter-lineage mechanisms represents an important direction for future work.

We found that host inflammatory responses were defined in two independent patterns. Production of core pro-inflammatory cytokines was comparatively similar across cells infected with different *P. aeruginosa*, consistent with pattern recognition through p38 mitogen activated protein kinase (MAPK) and NF-κB-dependent pathways (29–31). A variable tissue-remodeling response with significantly different levels of MMP-9 and VEGF production by cells infected with *P. aeruginosa* variants. MMP-9 and VEGF are associated with matrix degradation, neutrophil-driven airway injury, and airway remodeling in CF and bronchiectasis (32–34). The fact that a single variant such as P1V1 can shift the epithelium response along tissue-remodeling response, suggests that certain within-host adaptations act as switches for host responses between an acute clearance and a chronic remodeling program. If so, a key feature of pathogenesis in chronic infection may not be limited to the overall levels of inflammation, but the type of host response. Identifying *P. aeruginosa* ‘Variants of Concern’ within a patient’s bacterial population will be a critical step for accurately predicting disease progression and designing targeted therapeutics against chronic *P. aeruginosa*.

Beyond the global transcriptional signatures of *P. aeruginosa* in our infection model, our data highlight how populations from individual patients maintain highly stable transcription of key physiological switches, including *hfq*, *xcpT*, and *ssb*. Hfq is a global post-transcriptional regulator coordinating stress responses and virulence gene expression; XcpT is a component of the type II secretion machinery that exports much of the *P. aeruginosa* exoproteome; and Ssb is central to DNA replication and repair, consistent with the elevated *sulA* expression seen across clinical populations (35, 36). Stable expression of this set within populations, alongside divergence in individual variants, is more consistent with interaction-dependent modulation of collective behavior other than simply an average of members’ behavior. Whether these changes in collective behavior are due to regulatory cross-talk, division of labor among the variants, or frequency-dependent selection, is not resolved in this study.

Although previous studies report that *P. aeruginosa* transcriptional signatures are conserved across patients and independent of population structure (26–28), they address fundamentally different scales. Transcriptomes from whole expectorated sputum or from an explanted lung integrate *P. aeruginosa* across the airway, averaging spatially segregated lineages that may rarely interact directly. Our approach evaluated the micro-scale where lineages mingle, and functional interaction effects emerge. Considering the focal nature of airway injury in bronchiectasis and the spatial isolation of *P. aeruginosa* sub-populations (24, 37), clarifying dynamics at both the macro- and micro-scales is critical.

While constrained by a small participant cohort, our findings demonstrate a link between core cytokine output and ppFEV1, highlighting the impact of intra-strain diversity on host responses. We acknowledge that selecting one variant per morphotype does not reconstitute the entire population, and that this misses genetically distinct lineages with the same morphotype. Given that expectorated sputum samples only capture a fraction of *P. aeruginosa* load, leaving spatially segregated sub-populations in distant lung regions uncaptured, this remains a key limitation when defining collective bacterial behavior in chronic respiratory infections.

Taken together, our findings demonstrate that single bacterial isolates fail to capture the reality of chronic microbial pathogenesis. Functional interactions between co-existing sub-lineages actively reshape *P. aeruginosa* population physiology and host immune response, which makes the heterogeneous population, rather than individual isolates, a more relevant unit for the analysis of pathogenesis and disease progression. As CFTR modulator therapy changes the airway environment in which these populations persist, and as we broaden focus to chronic *P. aeruginosa* infections in non-CF bronchiectasis, determining how specific lineage compositions and frequencies activate damaging host remodeling pathways will be critical both for predicting disease progression and for engineering therapeutics that target collective population dynamics.

## Materials and methods

### Sputum sample collection and participants

We selected three adults with CF, aged 21–28 years, from a cohort at Emory University Hospital, Atlanta, with chronic *P. aeruginosa* infection for 10–15 years at the time of sputum collection, under Institutional Review Board protocols at Emory (IRB00042577) and Georgia State University (H23174). We collected and processed a single expectorated sputum sample from each participant as described previously (23). Briefly, we supplemented each sputum sample with 5 mL synthetic cystic fibrosis medium (SCFM) (38), homogenized by vortexing for 2 min, and centrifuged the homogenate for 4 min at ∼3,300 × *g*. We removed the supernatant, resuspended the pellet in phosphate-buffered saline, and inoculated 10-fold serial dilutions onto *Pseudomonas* isolation agar (PIA). We incubated plates at 37°C overnight and then at room temperature for up to 72 h. We collected all colonies from each sputum sample and stored them together as the whole population. For populations containing several morphotypes, we selected a representative variant of each morphotype. We amplified and Sanger sequenced the 16S rRNA gene to confirm *P. aeruginosa* identity for each selected variant before proceeding to whole-genome sequencing and phenotypic analysis.

### Colony morphology diversity in clinical populations

To assess heterogeneity in colony morphology, we used Congo red agar (1% agar, 1× M63 salts [3 g monobasic KH_2_PO_4_, 7 g K_2_HPO_4_, 2 g (NH_4_)_2_SO_4_, pH 7.4], 2.5 mM magnesium chloride, 0.4 mM calcium chloride, 0.1% casamino acids, 0.1% yeast extract, 40 mg/L Congo red, 100 µM ferrous ammonium sulfate, and 0.4% glycerol) (22, 39). We inoculated 10 µL of frozen whole *P. aeruginosa* populations into 5 mL SCFM and, after 6 h at 37°C with shaking at 200 rpm, plated 10 and 50 µL of culture onto Congo red agar. We incubated plates at 37°C for 24 h followed by 4 days at 22°C and archived a representative of each colony morphotype for genetic and phenotypic analysis.

### Primary CF airway epithelial cells and infection

We obtained differentiated primary CF airway epithelial cells at air–liquid interface (ALI) from the Cystic Fibrosis Biospecimen Repository (CFBR) core at Emory University. Briefly, P0 airway epithelial cells were resuspended in 20 mL E-ALI medium and counted. E-ALI medium contains insulin (5 µg/mL) and is supplemented with CaCl_2_ (1 mM), heparin (2 µg/mL), L-glutamine (2.5 mM), hydrocortisone (960 ng/mL), O-phosphoryl ethanolamine (0.5 µg/mL), bovine pituitary extract (20 µg/mL), and Mg^2+^ (0.5 µM). To differentiate the cells, 10^5^ cells per 6.5-mm well were seeded onto type IV collagen-coated trans-wells with 750 µL E-ALI in the basolateral chamber (40). Basolateral medium was replaced with fresh E-ALI and apical medium removed every 48 h to maintain ALI. We received fully differentiated cells from the CFBR core 14–21 days after transition to ALI.

For infection, we grew 10 µL of each whole mixed population or single variant in 3 mL SCFM at 37°C with shaking at 200 rpm for 6 h. We measured OD_600_ and adjusted each culture to deliver ≈10^3^ colony forming units/mL (CFU) in 100 µL SCFM2, followed by a further 2 h incubation at 37°C We washed the apical surface with 250 µL E-ALI, replaced the basolateral medium with fresh E-ALI, and infected by adding 100 µL of bacterial suspension in SCFM2 to the apical surface. Infected and uninfected control cells were incubated for 12 h at 37°C with 5% CO_2_; controls received an equal volume of SCFM2. After 12 h, we collected the basolateral medium for cytokine measurement and added up to 500 µL RNAlater (ThermoFisher) to the apical compartment for RNA extraction. We performed three independent experiments for each infection condition.

### Cytokine and chemokine measurement

We measured cytokines in cell-free basolateral supernatants (100 µL) by quantifying fluorescent intensity, using the Human Cytokine Array Q1 (RayBiotech) according to the manufacturer’s instructions. We determined the concentration of each cytokine attributable to infection by subtracting the value measured in cells treated with 100 µL SCFM2 alone for the same duration.

### RNA extraction and sequencing

We extracted RNA from two independent samples per condition stored in RNAlater. Samples were thawed on ice and RNAlater was removed by centrifugation. We resuspended pellets in RNase-free TE buffer containing 50 mg/mL lysozyme (v/v), incubated for 30 min at 37°C for enzymatic lysis, and added 1 mL TRI reagent (Sigma). We removed the aqueous phase and precipitated RNA with isopropanol. We depleted rRNA using NEBNext rRNA Depletion Kits (Human/Mouse/Rat and Bacteria) and fragmented RNA using the NEBNext Magnesium fragmentation module, following the manufacturer’s instructions (New England Biolabs). We prepared libraries with the NEBNext Small RNA Library Prep Kit and removed adapter dimers by size selection on a 5% TBE polyacrylamide gel (27). We sequenced libraries on an Illumina NextSeq 500 platform using 75-bp single-end runs at the Molecular Evolution Core, Georgia Institute of Technology.

### RNA-seq analysis

We assessed read quality with FastQC v0.11.8 (41)and used Cutadapt v2.6 (42) to remove Illumina adapters from the 3′ end, retaining reads of at least 22 bp. We mapped reads to *P. aeruginosa* PAO1 (GCF_000006765.1) using Bowtie 2 (43) and assigned mapped reads to PAO1 genes with featureCounts v2.0.1 (44) using the flags -s 1 (stranded) and -O (allowMultiOverlap). Downstream analyses used count data with variance-stabilizing transformation performed on all samples together in DESeq2 v1.28.1 (45) with blind = TRUE in R.

### Variance partitioning analysis

To quantify the contribution of population identity to gene expression variance, we used the variancePartition package (46). We partitioned total transcriptomic variance for each gene into experimental group (clinical populations versus PAO1; mixed populations versus individual variants) and unexplained residual variance, to identify expression patterns driving differences between groups rather than experimental noise.

### Statistical analysis

We performed all statistical analyses and data visualization in R v4.2.2 using the ggplot2, clusterProfiler, KEGGREST, and EnhancedVolcano packages.

## Data availability

RNA sequencing reads have been deposited in the NCBI Sequence Read Archive under BioProject PRJNA1510203.

## Acknowledgements

We thank all individuals with CF who provided sputum samples, and Arlene Stecenko and Katy Clemmer for assistance in acquiring sputum samples. Access to the Cystic Fibrosis Biospecimen Registry at the Emory Children’s Center for CF and Airways Disease Research was provided through the Emory University Pediatric CF Discovery Core under Institutional Review Board protocols at Emory (IRB00042577) and Georgia State University (H23174).

This work was supported by the Cystic Fibrosis Foundation through a fellowship to S.A. (AZIMI18F0) and grants to S.P.D. (DIGGLE18I0 and DIGGLE20G0), by CF@LANTA through a fellowship to S.A. (3206AXB), by Georgia State University Startup funds to S.A., and by the National Institutes of Health through grants to S.P.D. (R01AI153116; R56AI184449). Any opinions, findings, and conclusions or recommendations expressed in this material are those of the authors and do not necessarily reflect the views of the funding agencies.

## Author contributions

S.A. contributed to conceptualization, methodology development, investigation and writing. S.P.D. contributed to conceptualization and writing.

---

**Fig. S1.**
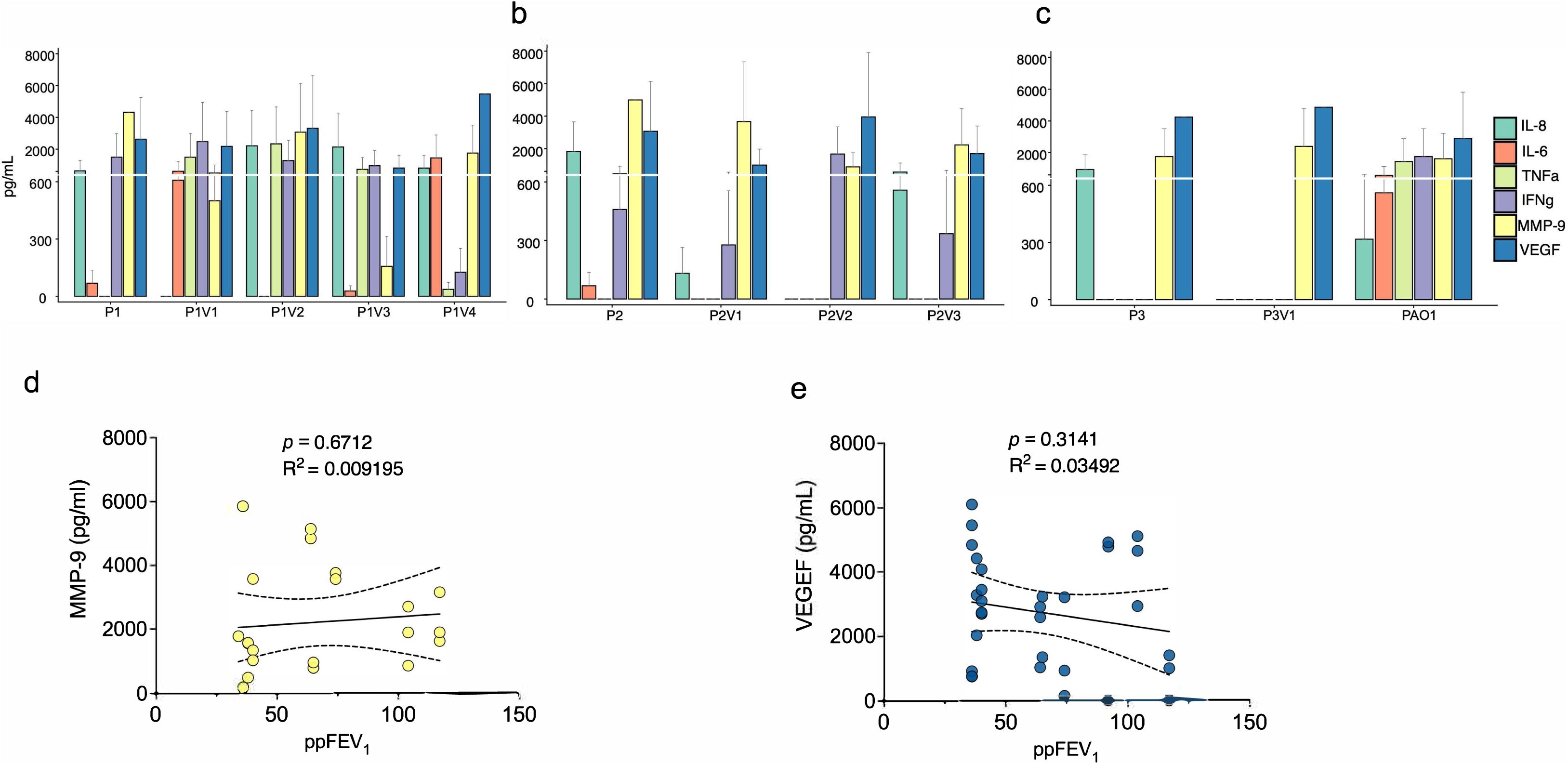
Individual variants induce distinct inflammatory and remodeling responses in CF-pAECs. (a–c) Pro-inflammatory cytokines (IL-6, IL-8, TNF-α, IFN-γ) and tissue-remodeling mediators (MMP-9, VEGF) produced by CF-pAECs following infection with mixed clinical *P. aeruginosa* populations, their clonal variants, and PAO1. Bars show mean concentration ± SEM for (a) P1 and variants P1V1–P1V4, (b) P2 and variants P2V1–P2V3, and (c) P3, variant P3V1, and PAO1. (d, e) Linear regression of participant ppFEV_1_ against secreted (d) MMP-9 (*R*^2^ = 0.009195, *P* = 0.6712) and (e) VEGF (*R*^2^ = 0.03492, *P* = 0.3141). Solid lines show best-fit linear regressions; dashed curves show 95% confidence intervals.

**Fig. S2.**
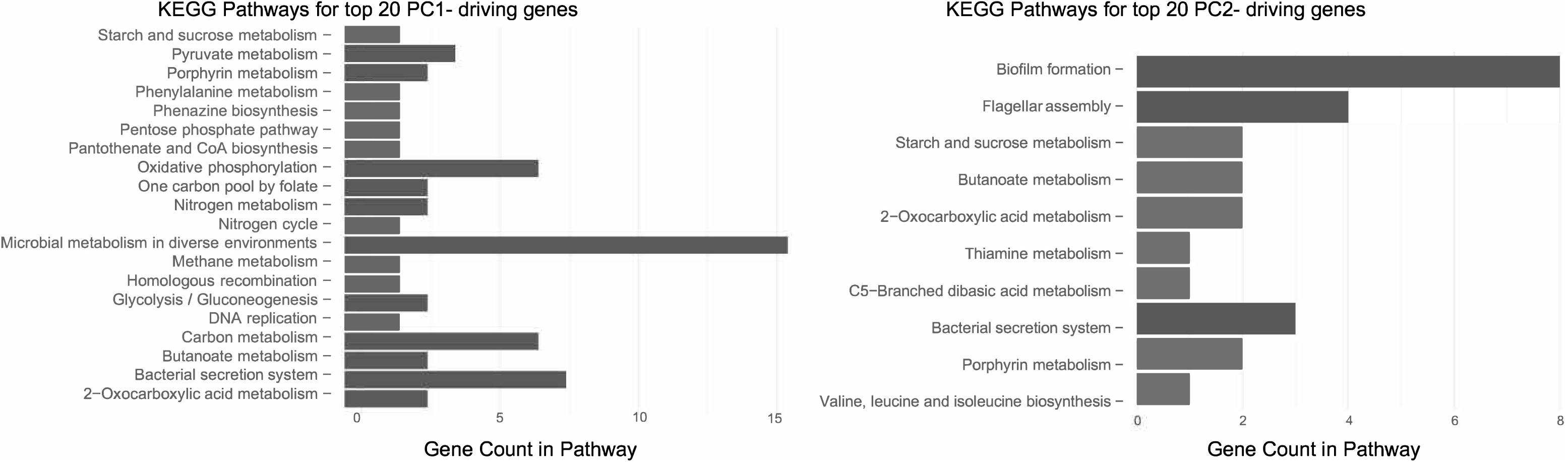
KEGG pathway enrichment for genes driving PC1 and PC2 in mixed *P. aeruginosa* populations. (a) Enrichment among the top 20 PC1-driving genes, including microbial metabolism in diverse environments, bacterial secretion systems, carbon metabolism, oxidative phosphorylation, and nitrogen metabolism. (b) Enrichment among the top 20 PC2-driving genes, including biofilm formation, flagellar assembly, bacterial secretion systems, and short-chain fatty acid and amino acid metabolism.

**Fig. S3.**
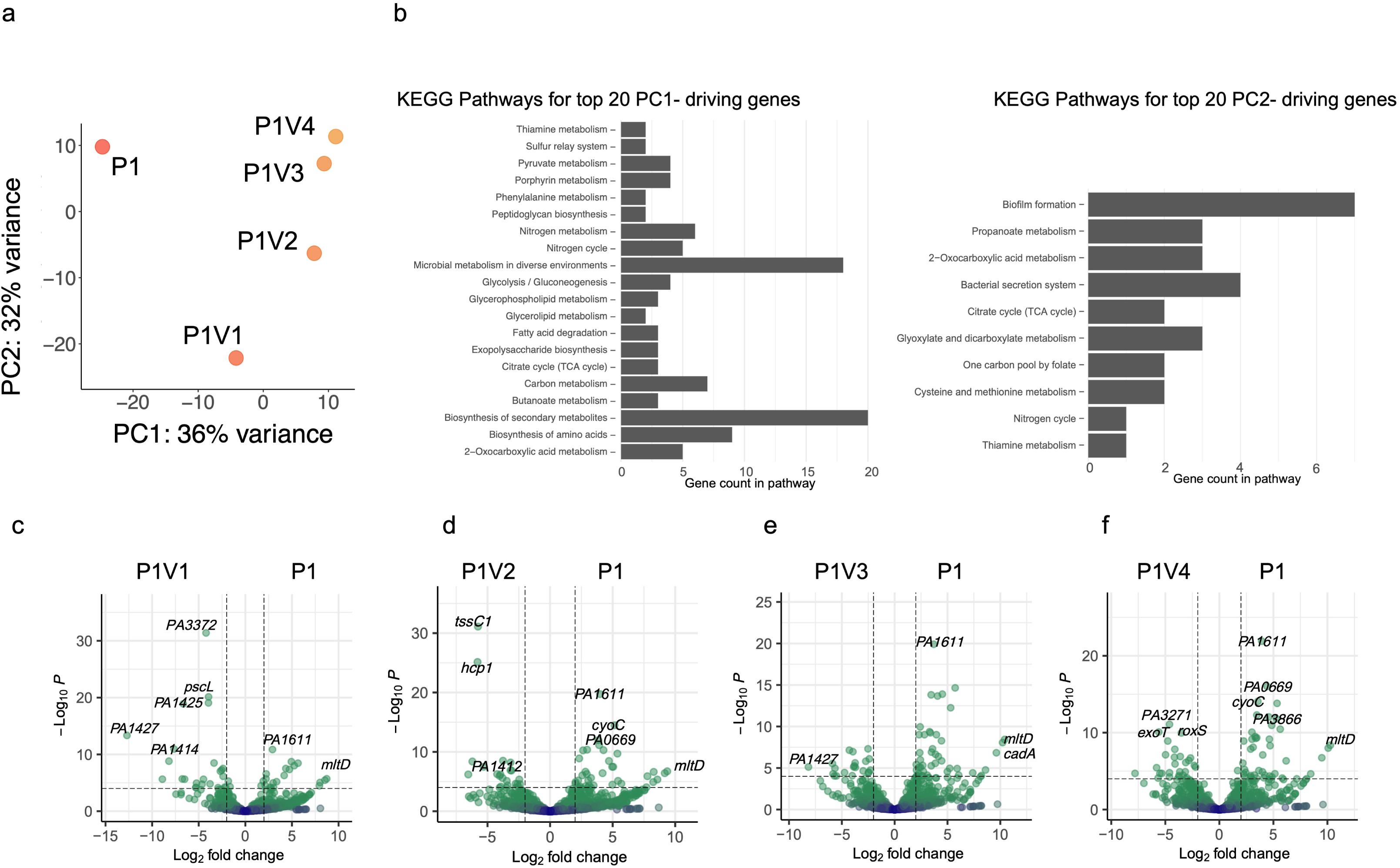
Transcriptomic divergence and pathway enrichment of clonal variants from population P1. (a) Global transcriptional differences between the P1 mixed population and variants P1V1–P1V4. PC1 (36%) and PC2 (32%) capture 68% of total variance. (b) PC1-driving genes are enriched for biosynthesis of secondary metabolites and carbon and nitrogen metabolism. (c) PC2-driving genes are enriched for biofilm formation, bacterial secretion systems, and the citrate (TCA) cycle. (d) Differential expression of P1V1 versus P1, showing downregulation of *PA3372*, *pscL*, *PA1425*, *PA1427*, and *PA1414*, and upregulation of *PA1611* and *mltD*. (e) Differential expression of P1V2 versus P1. (f) Differential expression of P1V3 versus P1, showing upregulation of *PA1611*, *mltD*, and *cadA* in the mixed P1 population. (g) Differential expression of P1V4 versus P1, showing differences in *exoT*, *roxS*, *PA3271*, *PA1611*, *PA0669*, *cyoC*, *PA3866*, and *mltD*. Dashed lines in volcano plots indicate cutoffs for log_2_ fold change (−2, 2) and adjusted *P* value (−log_10_ *P* > 3).

**Fig. S4.**
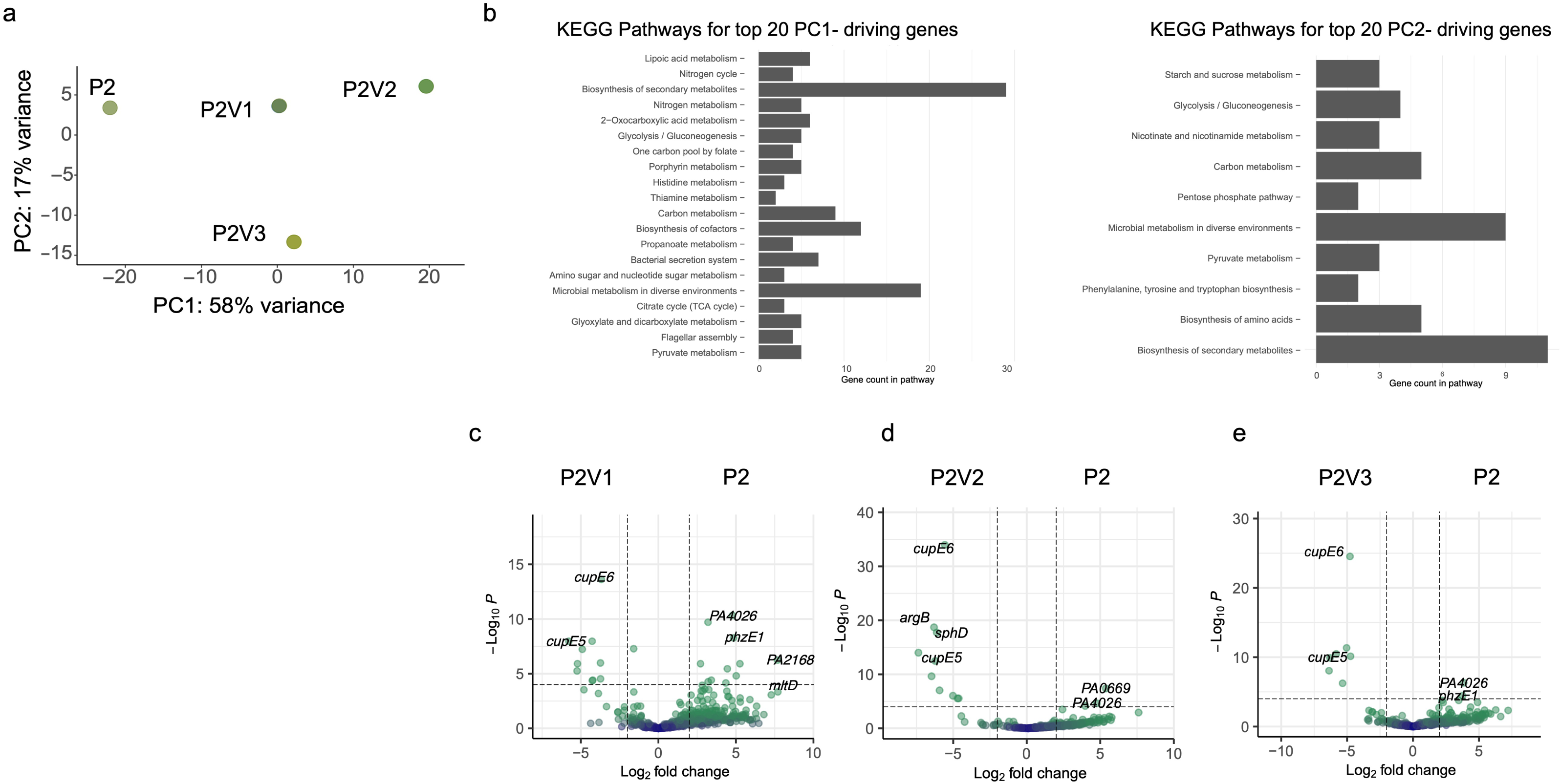
Transcriptomic divergence and pathway enrichment of clonal variants from population P2. (a) Global transcriptional differences between the P2 mixed population and variants P2V1–P2V3. PC1 (58%) and PC2 (17%) capture 75% of total variance, with P2V3 separating along PC2. (b) PC1-driving genes are enriched for biosynthesis of secondary metabolites, microbial metabolism in diverse environments, biosynthesis of cofactors, and carbon metabolism; PC2-driving genes are enriched for biosynthesis of secondary metabolites, microbial metabolism in diverse environments, carbon metabolism, and glycolysis/gluconeogenesis. (c) Differential expression of P2V1 versus P2, showing repression of the chaperone–usher fimbrial loci *cupE5* and *cupE6* in P2V1 and higher expression of *PA4026*, *phzE1*, *PA2168*, and *mltD* in the mixed population. (d) Differential expression of P2V2 versus P2, showing downregulation of *cupE6*, *argB*, *sphD*, and *cupE5*, and higher expression of *PA0669* and *PA4026* in the P2 population. (e) Differential expression of P2V3 versus P2, showing downregulation of *cupE5* and *cupE6* and upregulation of *phzE1*. Dashed lines indicate cutoffs for log_2_ fold change (−2, 2) and adjusted *P* value (−log_10_ *P* > 3).

**Fig. S5.**
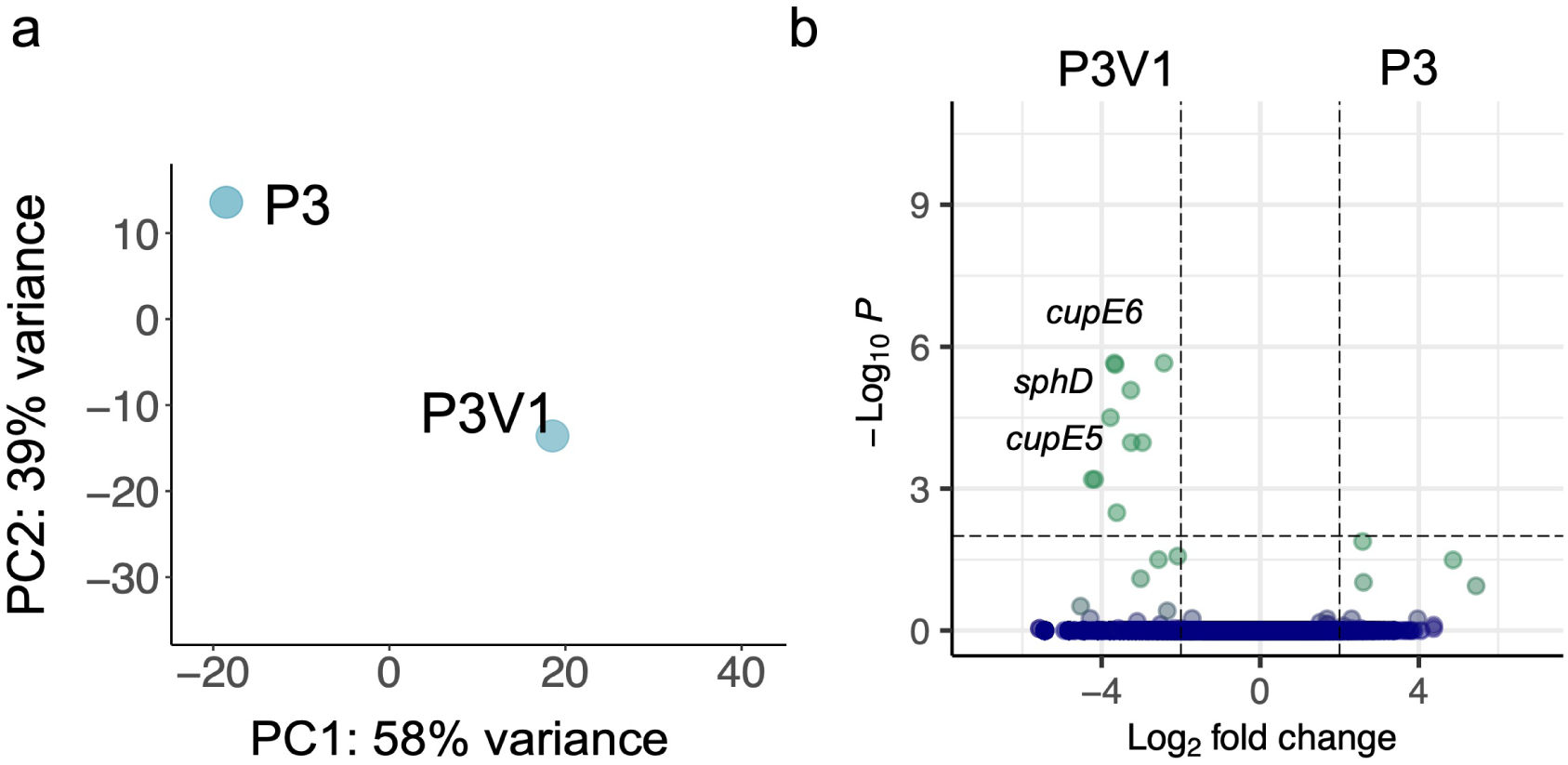
Transcriptomic divergence of the clonal variant from population P3. (a) Global transcriptional differences between the P3 mixed population and variant P3V1. PC1 (58%) and PC2 (39%) capture 97% of total variance, despite minimal population-level diversity in colony morphology. (b) *cupE5*, *cupE6*, and *sphD* are significantly downregulated in P3V1 relative to P3. Dashed lines indicate cutoffs for log_2_ fold change (−2, 2) and adjusted *P* value (−log_10_ *P* > 3).

## References

1. Azimi S, Lewin GR, Whiteley M. 2022. The biogeography of infection revisited. Nat Rev Microbiol doi:10.1038/s41579-022-00683-3.

2. Culyba MJ, Van Tyne D. 2021. Bacterial evolution during human infection: Adapt and live or adapt and die. PLoS Pathog 17:e1009872.

3. Didelot X, Walker AS, Peto TE, Crook DW, Wilson DJ. 2016. Within-host evolution of bacterial pathogens. Nat Rev Microbiol 14:150–62.

4. Dekker JP. 2024. Within-Host Evolution of Bacterial Pathogens in Acute and Chronic Infection. Annu Rev Pathol 19:203–226.

5. Garcia-Clemente M, de la Rosa D, Maiz L, Giron R, Blanco M, Olveira C, Canton R, Martinez-Garcia MA. 2020. Impact of *Pseudomonas aeruginosa* Infection on Patients with Chronic Inflammatory Airway Diseases. J Clin Med 9.

6. Greenwald MA, Wolfgang MC. 2022. The changing landscape of the cystic fibrosis lung environment: From the perspective of *Pseudomonas aeruginosa*. Curr Opin Pharmacol 65:102262.

7. Eklof J, Misiakou MA, Sivapalan P, Armbruster K, Browatzki A, Nielsen TL, Lapperre TS, Andreassen HF, Janner J, Ulrik CS, Gabrielaite M, Johansen HK, Jensen A, Nielsen TV, Hertz FB, Ghathian K, Calum H, Wilcke T, Seersholm N, Jensen JS, Marvig RL. 2022. Persistence and genetic adaptation of *Pseudomonas aeruginosa* in patients with chronic obstructive pulmonary disease. Clin Microbiol Infect 28:990–995.

8. Montgomery ST, Mall MA, Kicic A, Stick SM, Arest CF. 2017. Hypoxia and sterile inflammation in cystic fibrosis airways: mechanisms and potential therapies. Eur Respir J 49.

9. Courtney JM, Ennis M, Elborn JS. 2004. Cytokines and inflammatory mediators in cystic fibrosis. J Cyst Fibros 3:223–31.

10. Boucher RC. 2007. Airway surface dehydration in cystic fibrosis: pathogenesis and therapy. Annu Rev Med 58:157–70.

11. Elizur A, Cannon CL, Ferkol TW. 2008. Airway inflammation in cystic fibrosis. Chest 133:489–95.

12. Filkins LM, O’Toole GA. 2015. Cystic Fibrosis Lung Infections: Polymicrobial, Complex, and Hard to Treat. PLoS Pathog 11:e1005258.

13. Doring G, Parameswaran IG, Murphy TF. 2011. Differential adaptation of microbial pathogens to airways of patients with cystic fibrosis and chronic obstructive pulmonary disease. FEMS Microbiol Rev 35:124–46.

14. Rossi E, La Rosa R, Bartell JA, Marvig RL, Haagensen JAJ, Sommer LM, Molin S, Johansen HK. 2021. *Pseudomonas aeruginosa* adaptation and evolution in patients with cystic fibrosis. Nat Rev Microbiol 19:331–342.

15. Azimi S, Klementiev AD, Whiteley M, Diggle SP. 2020. Bacterial Quorum Sensing During Infection. Annu Rev Microbiol 74:201–219.

16. Fischer S, Klockgether J, Gonzalez Sorribes M, Dorda M, Wiehlmann L, Tummler B. 2021. Sequence diversity of the *Pseudomonas aeruginosa* population in loci that undergo microevolution in cystic fibrosis airways. Access Microbiol 3:000286.

17. Penesyan A, Kumar SS, Kamath K, Shathili AM, Venkatakrishnan V, Krisp C, Packer NH, Molloy MP, Paulsen IT. 2015. Genetically and Phenotypically Distinct *Pseudomonas aeruginosa* Cystic Fibrosis Isolates Share a Core Proteomic Signature. Plos One 10.

18. Higgs MG, Greenwald MA, Roca C, Macdonald JK, Sidders AE, Conlon BP, Wolfgang MC. 2025. Flagellar motility and the mucus environment influence aggregation-mediated antibiotic tolerance of *Pseudomonas aeruginosa* in chronic lung infection. mBio doi:10.1128/mbio.00831-25:e0083125.

19. Mould DL, Botelho NJ, Hogan DA. 2020. Intraspecies Signaling between Common Variants of *Pseudomonas aeruginosa* Increases Production of Quorum-Sensing-Controlled Virulence Factors. mBio 11.

20. Jorth P, Durfey S, Rezayat A, Garudathri J, Ratjen A, Staudinger BJ, Radey MC, Genatossio A, McNamara S, Cook DA, Aitken ML, Gibson RL, Yahr TL, Singh PK. 2021. Cystic Fibrosis Lung Function Decline after Within-Host Evolution Increases Virulence of Infecting *Pseudomonas aeruginosa*. Am J Respir Crit Care Med 203:637–640.

21. Saber MM, Donner J, Levade I, Acosta N, Parkins MD, Boyle B, Levesque RC, Nguyen D, Shapiro BJ. 2023. Single nucleotide variants in *Pseudomonas aeruginosa* populations from sputum correlate with baseline lung function and predict disease progression in individuals with cystic fibrosis. Microb Genom 9.

22. Azimi S, Roberts AEL, Peng S, Weitz JS, McNally A, Brown SP, Diggle SP. 2020. Allelic polymorphism shapes community function in evolving *Pseudomonas aeruginosa* populations. ISME J doi:10.1038/s41396-020-0652-0.

23. Vanderwoude J, Azimi S, Read TD, Diggle SP. 2024. The role of hypermutation and collateral sensitivity in antimicrobial resistance diversity of *Pseudomonas aeruginosa* populations in cystic fibrosis lung infection. mBio 15:e0310923.

24. Fraser HL, Moustafa DA, Goldberg JB, Azimi S. 2026. Whole-tissue imaging reveals intrastrain diversity shapes the spatial organization of *Pseudomonas aeruginosa* in a murine infection model. mSphere 11:e00657–25.

25. Azimi S, Thomas J, Cleland SE, Curtis JE, Goldberg JB, Diggle SP. 2021. O-Specific Antigen-Dependent Surface Hydrophobicity Mediates Aggregate Assembly Type in *Pseudomonas aeruginosa*. mBio doi:10.1128/mBio.00860-21:e0086021.

26. Cornforth DM, Diggle FL, Melvin JA, Bomberger JM, Whiteley M. 2020. Quantitative Framework for Model Evaluation in Microbiology Research Using *Pseudomonas aeruginosa* and Cystic Fibrosis Infection as a Test Case. mBio 11.

27. Cornforth DM, Dees JL, Ibberson CB, Huse HK, Mathiesen IH, Kirketerp-Moller K, Wolcott RD, Rumbaugh KP, Bjarnsholt T, Whiteley M. 2018. *Pseudomonas aeruginosa* transcriptome during human infection. Proceedings of the National Academy of Sciences of the United States of America 115:E5125–E5134.

28. Kordes A, Preusse M, Willger SD, Braubach P, Jonigk D, Haverich A, Warnecke G, Haussler S. 2019. Genetically diverse *Pseudomonas aeruginosa* populations display similar transcriptomic profiles in a cystic fibrosis explanted lung. Nat Commun 10:3397.

29. Malhotra S, Limoli DH, English AE, Parsek MR, Wozniak DJ. 2018. Mixed Communities of Mucoid and Nonmucoid *Pseudomonas aeruginosa* Exhibit Enhanced Resistance to Host Antimicrobials. MBio 9.

30. Coates MS, Alton E, Rapeport GW, Davies JC, Ito K. 2021. *Pseudomonas aeruginosa* induces p38MAP kinase-dependent IL-6 and CXCL8 release from bronchial epithelial cells via a Syk kinase pathway. PLoS One 16:e0246050.

31. Ribeiro CMP, Higgs MG, Muhlebach MS, Wolfgang MC, Borgatti M, Lampronti I, Cabrini G. 2023. Revisiting Host-Pathogen Interactions in Cystic Fibrosis Lungs in the Era of CFTR Modulators. Int J Mol Sci 24.

32. Fuentes-Zacarias P, Arzate-Castaneda DA, Sosa-Gonzalez I, Villeda-Gabriel G, Morales-Mendez I, Osorio-Caballero M, Helguera-Repetto AC, Diaz FN, Garcia-Lopez G, Flores-Herrera O, Arenas-Huertero F, Eslava-Campos C, Diaz-Ruiz O, Flores-Herrera H. 2021. *Pseudomonas aeruginosa* induces spatio-temporal secretion of IL-1beta, TNFalpha, proMMP-9, and reduction of epithelial E-cadherin in human alveolar epithelial type II (A549) cells. Acta Biochim Pol 68:207–215.

33. Gaggar A, Hector A, Bratcher PE, Mall MA, Griese M, Hartl D. 2011. The role of matrix metalloproteinases in cystic fibrosis lung disease. Eur Respir J 38:721–7.

34. Garratt LW, Sutanto EN, Ling KM, Looi K, Iosifidis T, Martinovich KM, Shaw NC, Kicic-Starcevich E, Knight DA, Ranganathan S, Stick SM, Kicic A, Australian Respiratory Early Surveillance Team for Cystic F. 2015. Matrix metalloproteinase activation by free neutrophil elastase contributes to bronchiectasis progression in early cystic fibrosis. Eur Respir J 46:384–94.

35. Stewart PS, Franklin MJ, Williamson KS, Folsom JP, Boegli L, James GA. 2015. Contribution of stress responses to antibiotic tolerance in *Pseudomonas aeruginosa* biofilms. Antimicrob Agents Chemother 59:3838–47.

36. Breidenstein EB, Bains M, Hancock RE. 2012. Involvement of the lon protease in the SOS response triggered by ciprofloxacin in *Pseudomonas aeruginosa* PAO1. Antimicrob Agents Chemother 56:2879–87.

37. Jorth P, Staudinger BJ, Wu X, Hisert KB, Hayden H, Garudathri J, Harding CL, Radey MC, Rezayat A, Bautista G, Berrington WR, Goddard AF, Zheng C, Angermeyer A, Brittnacher MJ, Kitzman J, Shendure J, Fligner CL, Mittler J, Aitken ML, Manoil C, Bruce JE, Yahr TL, Singh PK. 2015. Regional Isolation Drives Bacterial Diversification within Cystic Fibrosis Lungs. Cell Host Microbe 18:307–19.

38. Palmer KL, Aye LM, Whiteley M. 2007. Nutritional cues control *Pseudomonas aeruginosa* multicellular behavior in cystic fibrosis sputum. J Bacteriol 189:8079–87.

39. Mayer-Hamblett N, Rosenfeld M, Gibson RL, Ramsey BW, Kulasekara HD, Retsch-Bogart GZ, Morgan W, Wolter DJ, Pope CE, Houston LS, Kulasekara BR, Khan U, Burns JL, Miller SI, Hoffman LR. 2014. *Pseudomonas aeruginosa* in vitro phenotypes distinguish cystic fibrosis infection stages and outcomes. Am J Respir Crit Care Med 190:289–97.

40. Morgan R, Manfredi C, Easley KF, Watkins LD, Hunt WR, Goudy SL, Sorscher EJ, Koval M, Molina SA. 2022. A medium composition containing normal resting glucose that supports differentiation of primary human airway cells. Scientific Reports 12.

41. Andrews S. 2010. FastQC: A quality control tool for high throughput sequence data. https://www.bioinformatics.babraham.ac.uk/projects/fastqc/.

42. Martin M. 2011. Cutadapt removes adapter sequences from high-throughput sequencing reads. 2011 17:3.

43. Langmead B, Salzberg SL. 2012. Fast gapped-read alignment with Bowtie 2. Nat Methods 9:357–9.

44. Liao Y, Smyth GK, Shi W. 2014. featureCounts: an efficient general purpose program for assigning sequence reads to genomic features. Bioinformatics 30:923–30.

45. Love MI, Huber W, Anders S. 2014. Moderated estimation of fold change and dispersion for RNA-seq data with DESeq2. Genome Biol 15:550.

46. Hoffman GE, Schadt EE. 2016. variancePartition: interpreting drivers of variation in complex gene expression studies. BMC Bioinformatics 17:483.

